# Activation of MAIT cells plays a critical role in viral vector vaccine immunogenicity

**DOI:** 10.1101/661397

**Authors:** Nicholas M. Provine, Ali Amini, Lucy C. Garner, Christina Dold, Claire Hutchings, Michael E.B. FitzPatrick, Laura Silva Reyes, Senthil Chinnakannan, Blanche Oguti, Meriel Raymond, Stefania Capone, Antonella Folgori, Christine S. Rollier, Eleanor Barnes, Andrew J. Pollard, Paul Klenerman

## Abstract

Mucosal-associated invariant T (MAIT) cells can be activated by viruses through a cytokine-dependent mechanism, and thereby protect from lethal infection. Given this, we reasoned MAIT cells may have a critical role in the immunogenicity of replication-incompetent adenovirus vectors, which are novel and highly potent vaccine platforms. *In vitro*, ChAdOx1 (Chimpanzee Adenovirus Ox1) induced potent activation of MAIT cells. Activation required transduction of monocytes and plasmacytoid dendritic cells to produce IL-18 and IFN-α, respectively. IFN-α-induced monocyte-derived TNF-α was identified as a novel intermediate in this activation pathway, and activation required combinatorial signaling of all three cytokines. Furthermore, ChAdOx1-induced *in vivo* MAIT cell activation in both mice and human volunteers. Strikingly, MAIT cell activation was necessary *in vivo* for development of ChAdOx1-induced HCV-specific CD8 T cell responses. These findings define a novel role for MAIT cells in the immunogenicity of viral vector vaccines, with potential implications for future design.

**One sentence summary:** Robust immunogenicity of candidate adenovirus vaccine vectors requires the activation of unconventional T cells.

## Body

Mucosal-associated invariant T (MAIT) cells, an abundant T cell population in humans, bridge innate and adaptive immunity due to their ability to execute effector functions following cytokine stimulation in the absence of TCR signals(*1*). *In vivo*, MAIT cells can respond to viruses in this TCR-independent manner, and mediate protection against lethal infection via early amplification of local effector mechanisms(*2–4*). We reasoned that such focused activity could play a critical role in viral vaccine immunogenicity. Replication-incompetent adenovirus (Ad) vectors are novel and highly potent vaccine platforms for many human diseases(*5*). We therefore sought to determine if such vectors activate MAIT cells and if this activation impacts on vaccine immunogenicity.

Firstly, to determine if MAIT cells respond to Ad vectors, we stimulated human PBMCs for 24 h with increasing MOIs of Ad5 and ChAdOx1, two clinically-relevant vectors(*6, 7*). ChAdOx1 induced robust dose-dependent upregulation of IFN-γ, CD69, and granzyme B by MAIT cells (Fig. 1A-C; Fig. S1A-D). In contrast, Ad5 only weakly activated MAIT cells even at the maximum dose (Fig. 1A-C). Activation in response to Ad vectors was confirmed using the MR1/5-OP-RU tetramer to identify MAIT cells (Fig. S1E). Vδ2+ T cells share many characteristics with MAIT cells(*8, 9*), and showed analogous Ad vector-induced activation (Fig. S1A, S1F).

**Figure 1.**
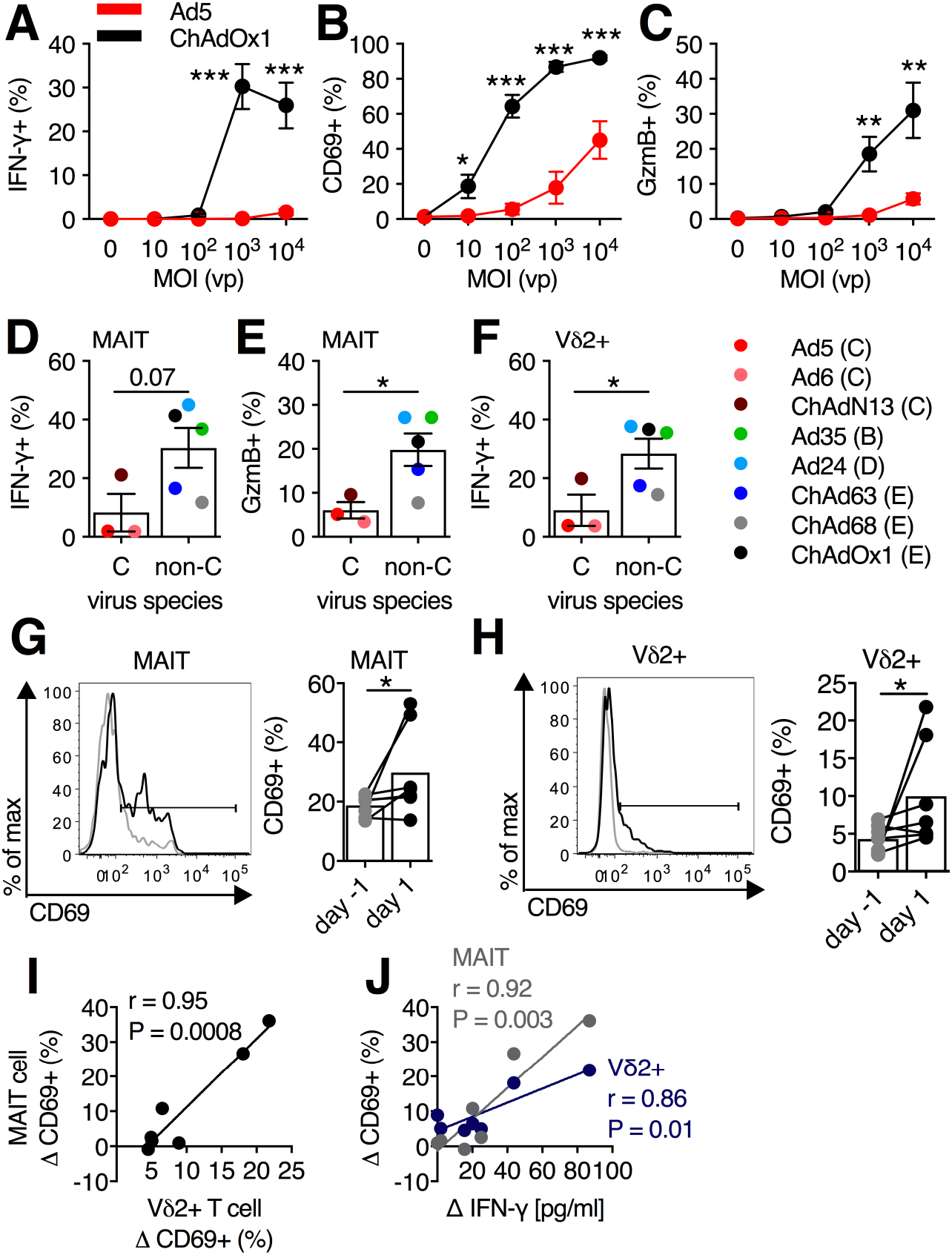
*In vitro* and *in vivo* activation of human MAIT and Vδ2+ T cells by adenovirus vectors. **(A-C)** PBMCs (N=9) were stimulated with Ad5 or ChAdOx1 at increasing MOIs (0 to 10^4^ vp), and IFN-γ **(A)**, CD69 **(B)**, and granzyme B (GzmB) **(C)** expression was measured on MAIT cells (CD161++Vα7.2+ T cells) after 24 h. **(D-F)** PBMCs were stimulated with MOI=10^3^ vp of the indicated vector (species in parentheses) for 24 h (N=5 per vector). Average IFN-γ **(D)** or GzmB **(E)** production by MAIT cells, and IFN-γ production by Vδ2+ T cells **(F)** in response to stimulation with the indicated vector. **(G-J)** Healthy human volunteers (N=7) were immunized with a 5×10^10^ vp dose of ChAdOx1 expressing a *N. meningitidis* group B antigen (MenB.1). Expression of CD69 on MAIT (MR1/5-OP-RU++ T cells) **(G)** and Vδ2+ T cells **(H)** in peripheral blood one day pre- and one day post-immunization. **(I)** Pearson correlation of change in CD69 expression on MAIT cells and Vδ2+ T cells following vaccination. **(J)** Pearson correlation of change in plasma IFN-γ level following vaccination with the change in expression of CD69 on MAIT cells and Vδ2+ T cells. *, P<0.05; **, P<0.01; ***, P<0.001. Unpaired T test **(A-F)** or Wilcoxon rank-sum test **(G,H)**. Symbols indicate average response of 5 donors for each vector **(D-F)** and individual donors **(G-J)**, and group mean (± SEM) are shown.

We tested a wider range of Ad vectors including three species C-derived vectors (weak innate inducers(*10–12*)): Ad5(*13*), Ad6(*13*), and ChAdN13 (unpublished), and five non-species C vectors (strong innate inducers(*10–12*)): Ad35 (B)(*13*), Ad24 (D)(*13*), ChAdOx1 (E) (*13, 14*), ChAd63 (E)(*13*), and ChAd68 (AdC68; E)(*15*). In response to stimulation with the various vectors there was a gradient of IFN-γ, CD69, and granzyme B production by MAIT and Vδ2+ T cells, which resulted in greater average activation by non-species C as compared to C vectors (Fig. 1D-F; Fig. S1G-I), consistent with the above reports of differential innate immune activation by these families of vectors.

We next determined if MAIT and Vδ2+ T cells are activated following administration of Ad vectors to humans. We analyzed the activation of MAIT and Vδ2+ T cells and plasma cytokine levels on day -1 and day 1 following immunization of humans with 5×10^10^ vp of a novel ChAdOx1-MenB.1 vaccine (Fig. S2A, S2B). We observed modest, but statistically-significant upregulation of CD69 on MAIT and Vδ2+ T cells one day following ChAdOx1 immunization (Fig. 1G, 1H), with no changes in cell frequency (Fig. S2C). The degree of MAIT and Vδ2+ T cell activation was highly correlated within individuals (Fig. 1I). Plasma cytokines/chemokines IFN-γ, IL-6, CCL-2, and TNF-α were induced following vaccination (Fig. S2D), consistent with data from non-human primates(*11*), and the degree of MAIT and Vδ2+ T cell activation was correlated with changes in these innate cytokines/chemokines (Fig. 1J; Fig. S2E).

The mechanism of Ad vector-induced activation of MAIT cells was next investigated. Ad5 and ChAdOx1 displayed similar abilities to transduce PBMCs (Fig. S3A), and HLA-DR+CD11c+CD19-CD3-monocytes/cDCs were the major transduced population (83-98% of GFP+ cells) (Fig. S3B, S3C). While Ad5 and ChAdOx1 both efficiently transduced monocytes/cDCs (Fig. 2A), Ad5 transduced only 1.5% of CD123+ pDCs compared with 17.4% of CD123+ pDCs transduced by ChAdOx1 (MOI=10^3^ vp) (Fig. 2A), consistent with a prior report of poor pDC transduction by Ad5(*12*).

**Figure 2.**
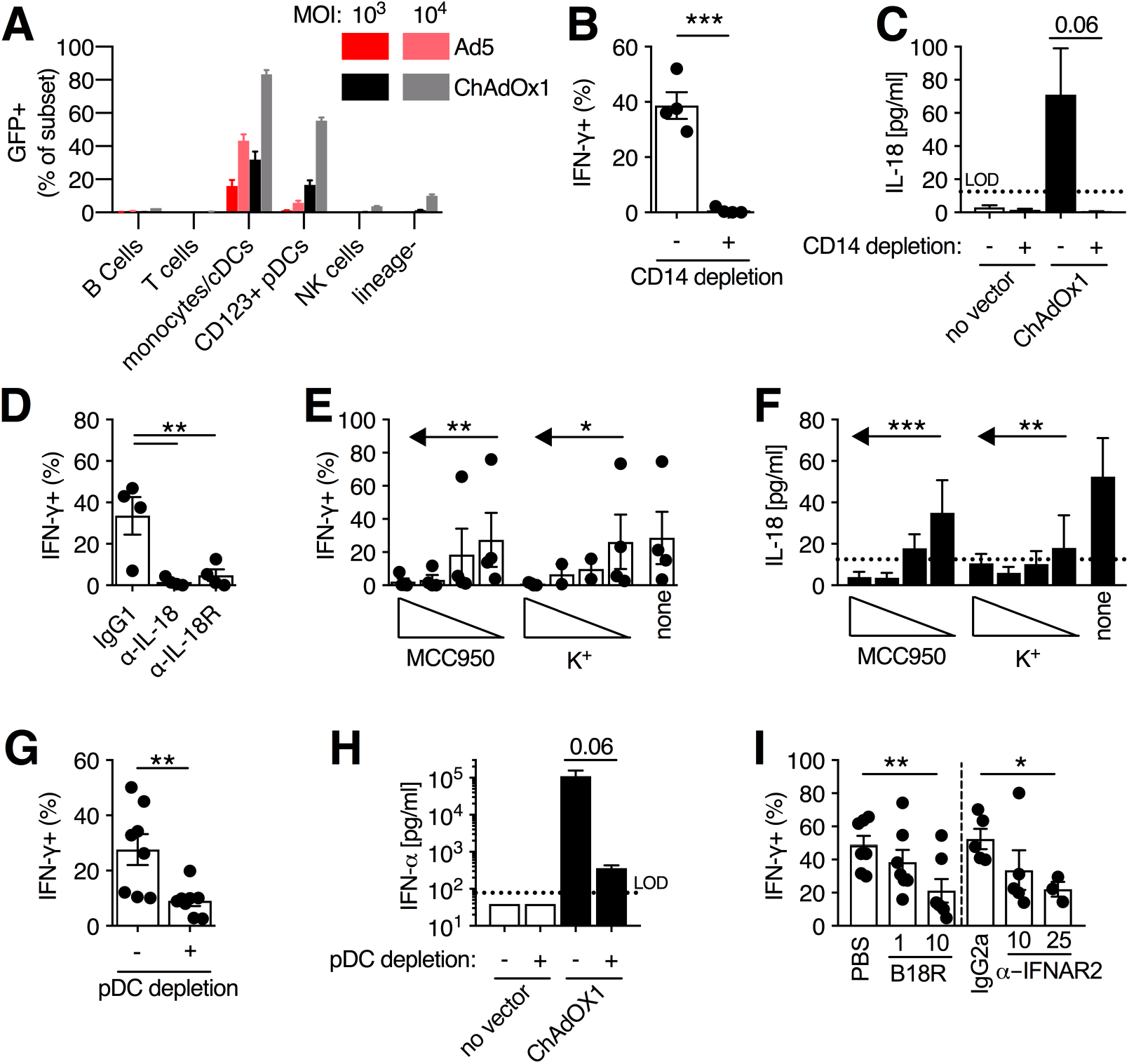
Activation of MAIT cells by adenovirus vectors requires monocyte-derived IL-18 and pDC-derived IFN-α. **(A)** Fresh human PBMCs (N=3) were stimulated for 24 h with either Ad5 or ChAdOx1 at MOI=10^3^ vp, and the fraction of each PBMC immune subset that was GFP+ was assessed. **(B)** PBMCs were depleted of CD14+ monocytes or left untreated as a control (N=4), and IFN-γ expression was measured on MAIT cells (CD161++Vα7.2+ T cells) after 24 h stimulation with ChAdOx1 (MOI=10^3^ vp). **(C)** Concentration of IL-18 in cell culture supernatants of whole PBMCs (N=5) or CD14-depleted PBMCs (N=4) 24 h after stimulation with ChAdOx1. **(D)** PBMCs (N=4) were treated with either anti-IL-18 or anti-IL-18R antibodies (10 μg/ml) immediately prior to stimulation with ChAdOx1 (MOI=10^3^ vp). IFN-γ production by MAIT cells was measured after 24 h. **(E,F)** The NLRP3 inhibitors (MCC950 [0.01-10 μM] and extracellular K+ [5-30 mM]) were added immediately prior to stimulation of PBMCs (N=4) with ChAdOx1. After 24 h, IFN-γ production by MAIT cells **(E)** and concentration of IL-18 in the cell culture supernatant **(F)** was assessed (N=4). **(G,H)** PBMCs were depleted of CD123+ pDCs or left untreated as a control (N=8), and IFN-γ expression on MAIT cells **(G)** or concentration of IFN-α in the cell culture supernatant **(H)** was measured after 24 h. **(I)** PBMCs were stimulated with ChAdOx1 (MOI=10^3^ vp) and B18R (1 or 10 μg/ml) or anti-IFNAR2 antibody (10 or 25 μg/ml) were added immediately prior to vector addition. IFN-γ expression was measured on MAIT-cells after 24 h. N=7 for B18R, and N=5 for anti-IFNAR2 antibody at 10 μg/ml and N=3 for 25 μg/ml. *, P<0.05; **, P<0.01; ***, P<0.001. Unpaired T test **(B,C,G,H)**, repeated-measures one-way ANOVA with Dunnett Correction **(D,I)**, repeated-measures one-way ANOVA with test for linear trend **(E,F)**. Symbols indicate individual donors, and mean ± SEM are shown.

Given their efficient transduction, we sought to determine the role of monocytes in Ad vector-induced activation of MAIT cells. Depletion of monocytes significantly reduced expression of IFN-γ, CD69, and granzyme B by MAIT cells following ChAdOx1 stimulation (Fig. 2B; Fig. S4A). Consistent with prior studies on viruses(*2, 3*), MAIT cell activation by Ad vectors was independent of TCR signaling (Fig. S4B) -- suggesting a cytokine-mediated activation process. Depletion of monocytes abolished IL-18 secretion following vector stimulation (Fig. 2C), and blockade of IL-18 signaling reduced MAIT cell IFN-γ, CD69, and granzyme B production (Fig. 2D; Fig. S4C). Blocking IL-12 reduced only IFN-γ production by MAIT cells (Fig. S4C), and blocking IL-15 had no effect. In contrast with ChAdOx1, Ad5 stimulation did not induce detectable levels of IL-18 or IL-12p70 (Fig S4D, S4E), consistent with the non-stimulatory nature of this vector. Direct inhibition of the Cathepsin B-NLRP3 inflammasome pathway(*16*) using four different pharmacologic approaches (Ca-074 Me, MCC950, elevated extracellular [K^+^], and Z-YVAD-FMK), significantly reduced expression of IFN-γ, CD69, and granzyme B by MAIT cells (Fig. 2E; Fig. S5A-C), and production of IL-18 following ChAdOx1 stimulation (Fig. 2F; Fig. S5D), similar to prior data examining IL-1β(*17, 18*). This effect was not due to altered transduction of PBMCs by ChAdOx1 (Fig. S5E).

Given the differential transduction of pDCs, the role of these cells in Ad vector-mediated activation of MAIT cells was investigated. Depletion of CD123+ pDCs resulted in a significant 67% reduction in IFN-γ production by MAIT cells (Fig. 2G), and reduced IFN-α levels by >99% following ChAdOx1 stimulation (Fig. 2H). Inhibition of type I interferon signaling reduced IFN-γ production by MAIT cells by 56-58% (Fig. 2I). Compared with ChAdOx1, Ad5 induced negligible amounts of IFN-α (Fig. S6A, S6B), consistent with previous reports(*11, 12*).

We envisaged a model where monocyte-derived IL-18 and pDC-derived IFN-α were the minimal factors required to activate MAIT cells in response to ChAdOx1 stimulation. However, while IFN-α/β + IL-18 induced MAIT cell IFN-γ in a PBMC culture, this was not seen using isolated MAIT cells (Fig. 3A), despite these cytokines upregulating CD69 on isolated MAIT cells (Fig. S7A). Depletion of monocytes from PBMCs reduced MAIT cell IFN-γ production following IFN-α + IL-18 stimulation (Fig. S7B), and addition of monocytes rescued this (Fig. 3B), indicating a monocyte-derived, IFN-α-dependent factor. The stimulatory factor was secreted, as either conditioned supernatant from IFN-α-treated monocytes (combined with IL-18), or provision of PBMCs across a transwell, significantly rescued IFN-γ production by isolated MAIT cells (Fig. 3C; Fig. S7C). IFN-α-stimulated monocytes secreted multiple interferon-responsive chemokines (e.g. MCP-1/CCL2), as well as TNF-α (Fig. 3D; Fig. S7D). Addition of recombinant TNF-α or an anti-TNFR2 agonist to IFN-α + IL-18-stimulated isolated MAIT cells increased IFN-γ production by >300% (from 4% to 16.5% and 17.6%, respectively; Fig. 3E; Fig. S7E). We confirmed the critical role of TNF-α as the presence of anti-TNF-α antibody (adalimumab) during IFN-α + IL-18 stimulation of PBMCs inhibited IFN-γ production by MAIT cells (Fig. S7F). The stimulatory capacity of supernatant from IFN-α-conditioned monocytes was also inhibited by the presence of adalimumab (Fig. S7G). TNF-α blockae using either adalimumab or recombinant TNFR2-Fc fusion protein (etanercept), but not a control anti-α4β7 antibody (vedolizumab), inhibited IFN-γ production by MAIT cells in response to ChAdOx1 (Fig. 3F; Fig. S7H). Depletion of monocytes reduced ChAdOx1-induced TNF-α production by 94% (Fig. 3G). Furthermore, Ad5 induced minimal TNF-α as compared with ChAdOx1 (Fig. S7I), consistent with the differential capacity of these two vectors to stimulate IFN-α production by pDCs (Fig. S6A).

**Figure 3.**
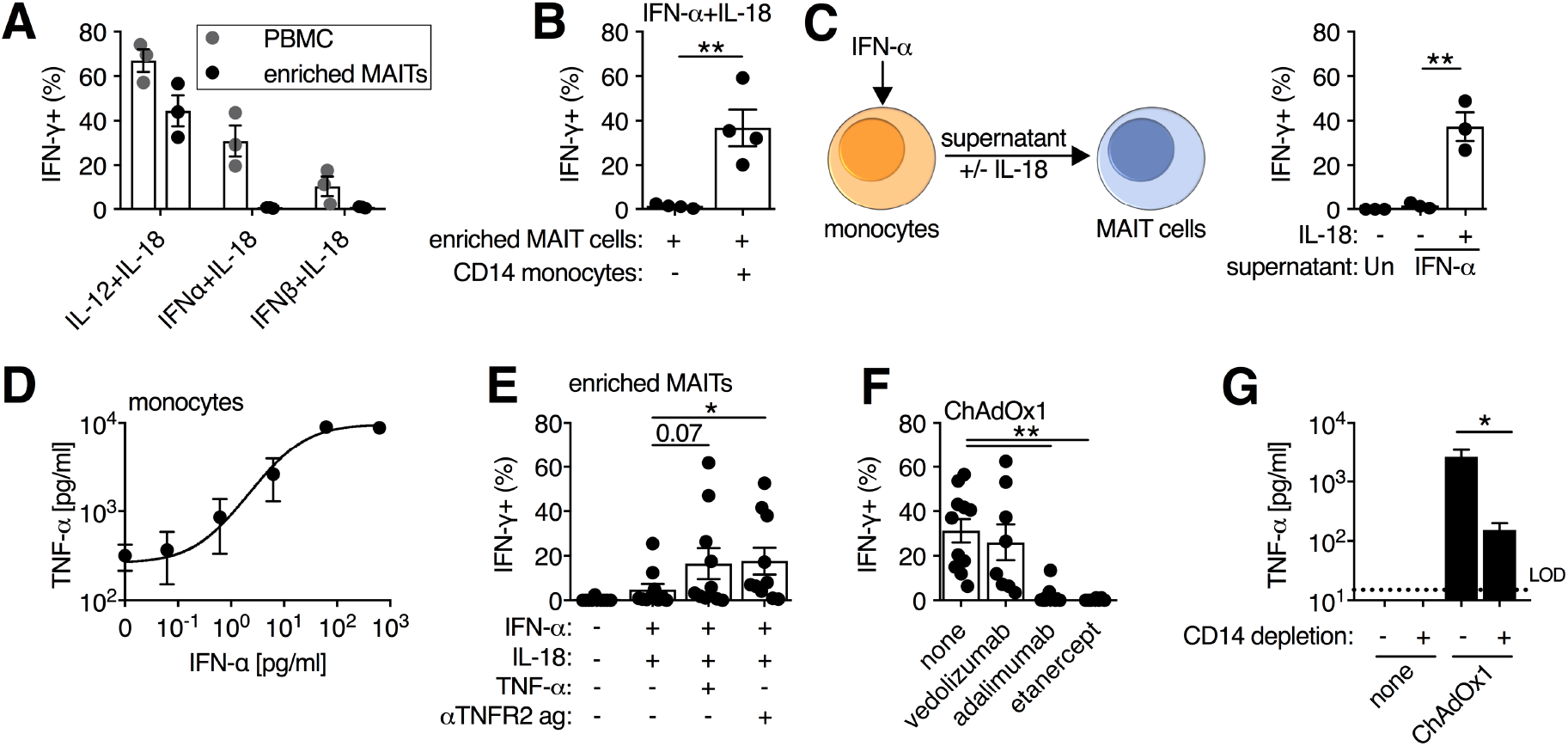
IFN-α acts directly and indirectly through the induction of TNF-α to activate MAIT cells. **(A)** Unfractionated PBMCs or enriched MAIT cells (positive selection by CD8 MicroBeads) were stimulated for 24 h with the indicated cytokines (50 ng/ml), and IFN-γ expression was measured on MAIT cells (CD161++Vα7.2+ T cells) after 24 h (N=3). **(B)** Enriched MAIT cells with or without CD14+ monocytes (positive selection by CD14 MicroBeads) were stimulated with IFN-α and IL-18 (50 ng/ml). IFN-γ production by MAIT cells was measured after 24 h (N=4). **(C)** Purified monocytes (N=3) were stimulated for with IFN-α (50 ng/ml), or left untreated, and after 24 h supernatants were transferred to autologous enriched MAIT cells with or without the addition of IL-18 (50 ng/ml). IFN-γ production by MAIT cells was measured after 24 h. **(D)** CD14-purified monocytes (N=3) were stimulated with increasing concentrations of IFN-α and TNF-α concentration in the cell culture supernatant was measured after 24 h. **(E)** Enriched MAIT cells (N=10) were stimulated with IFN-α and IL-18 ± TNF-α (50 ng/ml) or anti-TNFR2 agonist antibody (2.5 μg/ml), and IFN-γ production by MAIT cells was measured at 24 h. **(F)** PBMCs were stimulated with ChAdOx1 (MOI=10^3^ vp) and vedolizumab (anti-α4β7 integrin antibody, N=8), adalimumab (anti-TNF-α antibody, N=11), or etanercept (TNFR2-Fc fusion protein, N=8) (10 μg/ml) were added immediately prior to vector addition. IFN-γ production by MAIT cells was measured after 24 h. **(G)** Concentration of TNF-α in cell culture supernatants of whole PBMCs or CD14-depleted PBMCs 24 h after stimulation with ChAdOx1 (MOI=10^3^ vp; N=4). *, P<0.05; **, P<0.01. Unpaired T test **(B,C,G)**, repeated-measures one-way ANOVA with Dunnett Correction **(E,F)**. Symbols indicate individual donors, and mean ± SEM are shown.

Vδ2+ T cells were activated by Ad vectors through similar mechanisms (Fig. S8A-G). Compiling the data, the activation of innate-like T cells in response to Ad vectors requires the concerted action of IFN-α, TNF-α, and IL-18 (Fig. S9). These data extend prior reports of IFN-α-dependent activation of MAIT and Vδ2+ T cells by viruses(*3, 19*), by identifying a novel role for TNF-α as a necessary critical intermediary in this signaling pathway.

We next sought to determine the impact of MAIT cell activation on the induction of conventional T cell responses by ChAdOx1 immunization. C57BL/6J mice were immunized intramuscularly with ChAdOx1 or Ad5 at 10^8^ IU, and MAIT cell activation in the spleen, liver, and inguinal LNs was measured on day 1 (Fig. S10A-C). ChAdOx1 induced substantial upregulation of CD69 and granzyme B on MAIT cells in the inguinal LNs, and to a lesser degree in the liver (Fig. 4A, 4B). Ad5 induced significantly less expression of CD69 and granzyme B. In mice, iNKT cells are the most abundant innate-like T cell population(*20*), and CD69 and granzyme B were also significantly upregulated on iNKT cells following ChAdOx1 immunization, with Ad5 inducing less activation (Fig. S10D, S10E). These findings validate the use of a mouse model, as these data recapitulate the (differential) activation of MAIT cells by Ad vectors.

**Figure 4.**
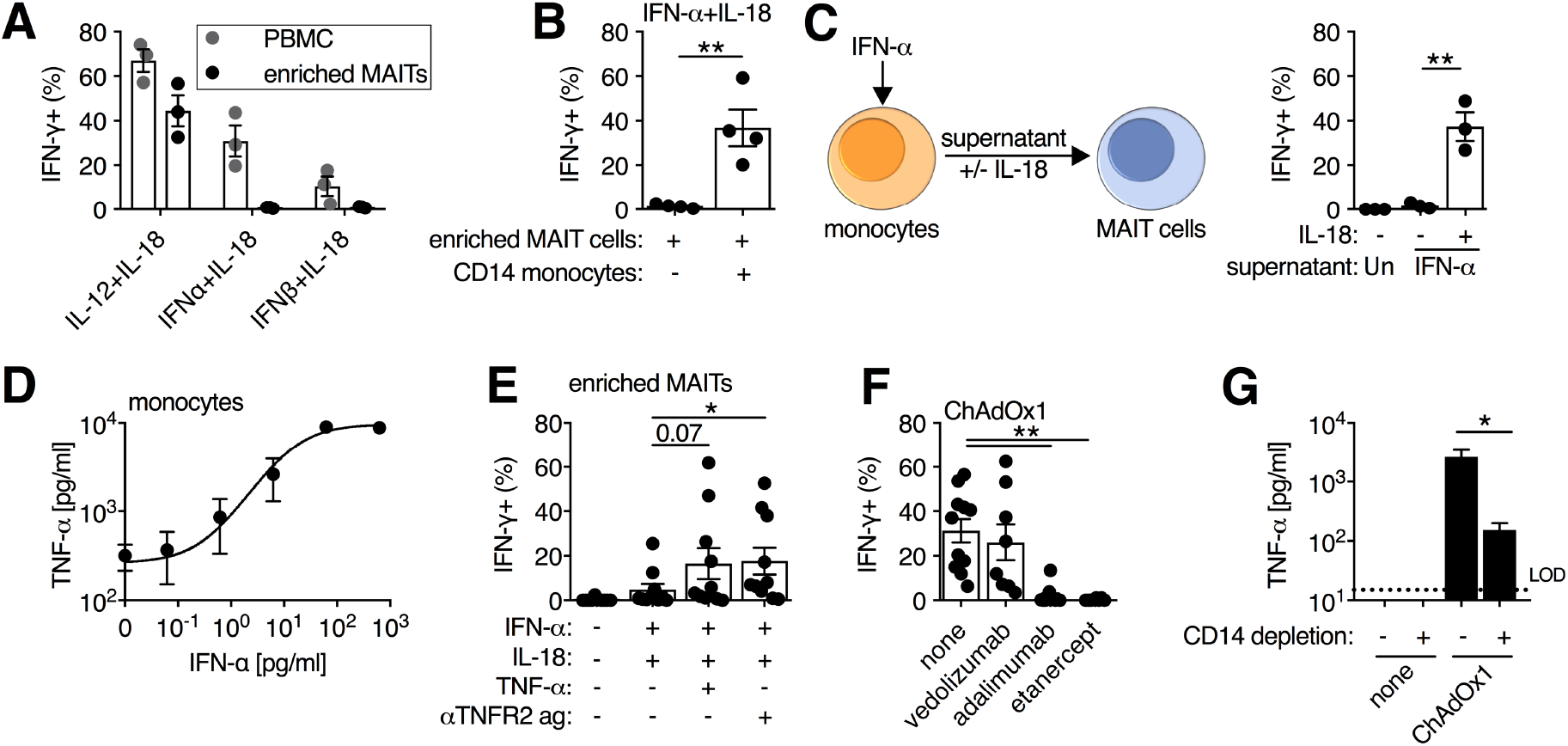
MAIT cell-deficient mice have impaired vaccine-induced CD8 T cell responses following ChAdOx1 immunization. **(A,B)** C57BL/6J mice (N=6 per group) were immunized intramuscularly with 10^8^ IU of Ad5 or ChAdOx1 expressing GFP, and one day post-immunization, expression of CD69 **(A)** and granzyme B (GzmB) **(B)** was measured on MAIT cells (MR1/5-OP-RU+ T cells) isolated from inguinal LNs and liver. Data are representative of two independent experiments. (C-G) C57BL/6J (N=12) or MR1 KO (N=9) mice were immunized intramuscularly with 10^8^ IU of ChAdOx1 expressing HCV-GT1-6_D_TM-Ii+L transgene, and on day 16 post-immunization splenocytes were collected. Representative flow cytometry plots **(C)** and group averages of IFN-γ production **(D)**, TNF-α production **(E)**, or dual production of IFN-γ an TNF-α **(F)** by CD8 T cells following 5 h restimulation with an overlapping peptide pool of the HCV genotype 1b proteome. **(G)** Representative flow cytometry plots and group averages of CD107a expression on cytokine-producing CD8 T cells following peptide restimulation. *, P<0.05; **, P<0.01; ***, P<0.001. One-way ANOVA with Sidak correction for multiple comparisons **(A,B)**, Unpaired T test **(D-G)**. Symbols indicate individual animals, and mean ± SEM are shown.

Having validated the model, we next addressed the role of MAIT cells in immunogenicity of Ad vector vaccines. Wildtype (WT) C57BL/6J and MR1 KO mice (Fig. S10F-H)(*21*) were immunized intramuscularly with 10^8^ IU of ChAdOx1 expressing an optimized invariant chain-linked HCV antigen(*22, 23*), and HCV-specific immune responses were measured on day 16 post-immunization. Following vaccination, MR1 KO mice had significantly reduced frequencies of CD8 T cells that produced IFN-γ, TNF-α, or both IFN-γ and TNF-α in response to HCV peptides, as compared with WT mice (Fig. 4C-F). This functional defect appeared specific to the CD8 T cell compartment, as there was no significant reduction in the frequency of HCV-specific CD4 T cells following vaccination of MR1 KO mice (Fig. S10I). HCV-specific CD8 T cells from MR1 KO mice also showed reduced degranulation, as measured by CD107a (Fig. 4G), and these cells displayed impaired differentiation towards KLRG1+ effector cells (Fig. S10J).

In summary, MAIT cells are capable of sensing the diversity of the Ad vector-induced innate immune activation landscape (e.g. IFN-α, TNF-α, IL-18) and can integrate these signals to augment vaccine-induced adaptive immune responses. The blend of signals required to maximally trigger MAIT cells uncovered here includes a novel and critical pathway via IFN-dependent TNF-α release, relying on cross-talk between two distinct populations of transduced cells, and varying between adenovirus serotypes. This full integration process is required for robust IFN-γ production, which has been shown to be critical for MAIT cell-mediated protection from viral infection(*4*).

This non-redundant role for MAIT cells places them in a critical bridging position between innate and adaptive immunity, despite many potentially shared functions with other innate-like populations(*9*). These data, coupled with studies in the lung(4, 24, 25), support an emerging model that MAIT cells can function to orchestrate early events in T cell-mediated immunity. It is striking that activation of MAIT cells – an abundant human innate-like population – is tightly and mechanistically linked to the immunogenicity of adenovirus vectors, which have emerged as a potent platform for T cell immunogenicity in human clinical trials(*26, 27*). This knowledge can be harnessed to further improve the design and development of these – and potentially other – vaccines against infections and cancer.

## Acknowledgements

We would like to thank: Stephanie Slevin and Helen Ferry for assistance with flow cytometry and panel design, Carl-Philipp Hackstein and Christian Willberg for critical discussions, Mariolina Salio and Vincenzo Cerundolo for provision of MR1 KO mice, Marilù Esposito, Hussein Al-Mossawi, Lian Ni Lee, and Timothy Donnison for provision of reagents, the NIH Tetramer Facility for provision of MR1 and CD1d tetramers, and all of the volunteers for participation in the trial and donation of blood samples.

## Author contributions

NMP and PK designed the project.

NMP, CD, CSR, EB, AJP, and PK designed the experiments.

NMP, AA, LCG, CD, CH, MEBF, LSR performed the experiments.

LSR, SC, BO, MR, SC, AF, CR, EB, and AJP provided samples and reagents.

All authors contributed to the writing and editing of the manuscript.

## Data Availability

All primary data available upon request.

## Supplementary Material

### Materials and Methods

#### Vectors and viruses

E1/E3-deleted replication-incompetent recombinant Ad5-GFP (VP:PFU ratio batch 1: 34, batch 2: 15, batch 3: 21), ChAdOx1-GFP (VP:PFU ratio batch 1: 118, batch 2: 13, batch 3: 78, batch 4: 73), ChAdOx1-HCV-GT1-6_D_TM-Ii+L(*22*) (VP:PFU ratio 95), and ChAd63-GFP (VP:PFU ratio 107) adenovirus vectors were produced by the Jenner Institute Viral Vector Core Facility at the University of Oxford, as previously described(*14*). ChAdOx1-MenB.1 (VP:PFU ratio 96) was produced at the Clinical Biomanufacturing Facility at the University of Oxford as previously described(*28*). E1/E3-deleted replication-incompetent recombinant Ad6 (VP:PFU ratio 95), ChAdN13 (VP:PFU ratio not calculated), Ad24 (VP:PFU ratio not calculated), Ad35 (VP:PFU ratio 124), and ChAd68 (AdC68; VP:PFU ratio 100) adenovirus vectors were produced by Nouscom, SRL (Rome, Italy), as previously described(*29*). Briefly, vectors were propagated in HEK293, except for ChAdOx1-MenB.1 which was propagated in PER.C6 cells, and isolated by CsCl_2_ ultracentrifugation.

#### Human PBMCs and isolation of cell populations

Fresh blood from healthy human volunteers was collected in EDTA-coated Vacutainer tubes (BD Biosciences) under the “Gastrointestinal Illness in Oxford: prospective cohort for outcomes, treatment, predictors and biobanking” (Ref: 11/YH/0020) ethics, or blood from anonymized healthy donors was collected from the NHS Blood and Transplant Service. Peripheral blood mononuclear cells (PBMCs) were isolated by density gradient centrifugation, as previously described(*8*). Briefly, PBS was used to dilute blood prior to layering over Lymphoprep (Axis-Shield or STEMCELL Technologies). Samples were centrifuged at 973 *g* for 30 minutes and allowed to decelerate without the brake. The PBMC layer was collected and washed once in R10 media [RPMI-1640 (Lonza) + 10% FBS (Sigma Aldrich) + 1% penicillin/streptomycin (Sigma-Aldrich)]. Red blood cells were lysed by incubation of the cell pellet in an 1x Ammonium-Chloride-Potassium (ACK) solution for <5 min. Cells were washed again in R10, and either used immediately or stored in liquid nitrogen.

CD8 MicroBeads (Miltenyi Biotec) were used to generate enriched MAIT-cell populations. CD14 MicroBeads and CD123 MicroBeads (Miltenyi Biotec) were used to deplete monocytes and plasmacytoid DCs (pDCs), respectively. All kits were used as per the manufacturer’s instructions.

#### Vaccinated human volunteers

PBMCs and plasma were collected from healthy volunteers aged 18-50 enrolled in the clinical trial ISRCTN trial number: ISRCTN46336916. Briefly, volunteers received a homologous prime-boost of 5×10^10^ vp of ChAdOx1-MenB.1 at a 6-month interval. Samples were collected prior to the second immunization and 1 day after.

#### Mice and tissue processing

JAX™ C57BL/6J mice (aged 6-10 weeks) were purchased from Charles River. MR1 KO mice(*21*) (kindly provided by Mariolina Salio and Vincenzo Cerundolo, University of Oxford) were bred in house and used at 6-10 weeks of age. Sex and age were matched between groups. All animals were housed in specific pathogen-free conditions at the Biomedical Services Building (University of Oxford) or the Wellcome Centre for Human Genetics (University of Oxford). All work was performed under UK Home Office license PPL 30/3293 or 30/3386 in accordance with the UK Animal (Scientific Procedures) Act 1986. All work was performed by trained and licensed individuals.

Animals were immunized intramuscularly in the hind legs with 10^8^ infectious units (IU) of Ad5-GFP, ChAdOx1-GFP, or ChAdOx1-HCV-GT1-6_D_TM-Ii+L, as previously described(*22*). Spleen and lymph nodes were processed as described previously(*30*). Briefly, tissue was dissociated through a 70 μm filter and washed with R10 media. Red blood cells were lysed, as needed, with 1x ACK solution for <5 min, and cells were washed an additional time with R10 media before downstream applications. Liver tissue was processed as described previously(*31*). Briefly, liver tissue was ground through a 70 μm filter, and washed once with R10 media. Liver mononuclear cells (MNCs) were isolated on a 35%-70% discontinuous Percoll (GE Healthcare) gradient by centrifugation at 741 *g* for 20 min and deceleration was without the brake. The MNC layer was collected, samples were washed once with R10 media, and residual red blood cells were lysed with 1x ACK solution for <5 min. After a final wash with R10 media, liver MNCs were used for downstream applications.

#### Genotyping

Genotyping of MR1 KO mice was performed by PCR and gel electrophoresis, as previously described(*21*). DNA was extracted from splenocytes using the Qiagen DNeasy Blood and Tissue Extraction kit per the manufacturer’s instructions. For testing of wildtype *Mr1* the MR1 5’ 8763-8783 (AGC TGA AGT CTT TCC AGA TCG) and MR1 9188-9168 rev (ACA GTC ACA CCT GAG TGG TTG) primers were used. For mutant *Mr1* the MR1 5’ 8763-8783 (AGC TGA AGT CTT TCC AGA TCG) and MR1 10451-10431 (GAT TCT GTG AAC CCT TGC TTC) primers were used. Primers (Sigma-Aldrich) were used at 10 μM and QuantiFast SYBR Green Mastermix (Qiagen) was used. Thermocycler conditions were: Step 1: 95 °C for 5 min, Step 2: 94 °C for 30 sec, Step 3: 60 °C for 30 sec, Step 4: 72 °C for 30 sec, Step 5: repeat Step 2-4 35x, Step 6: 72 °C for 5 Min, Step 7: hold at 22 °C. PCR products were run on a 2% Agarose gel and imaged on a GelDoc-It (UVP Imaging).

#### *In vitro* MAIT and Vδ2+ T cell stimulation assays

For *in vitro* stimulation of human PBMCs with Ad vectors, fresh PBMCs were used. For *in vitro* stimulation of human PBMCs with cytokines, fresh or freeze-thawed PBMCs were used with equivalent outcomes. For Ad vector stimulations, a previously described protocol(*11*) was used with slight modifications. 10^6^ whole PBMCs or cell subset-depleted PBMCs were added to a 96 well U-bottom plate. Ad-vectors were added at an MOI of 10^3^ vp, unless indicated otherwise. If applicable, inhibitory compounds or recombinant cytokines were added immediately prior to addition of the vector. Samples were mixed and incubated at 37 °C in 5% CO_2_.

For cytokine stimulations, a previously-described protocol(*1*) was used with slight modifications. Briefly, 10^6^ whole PBMCs or cell subset-depleted PBMCs were added to a 96 well U-bottom plate, and for isolated CD14+ monocytes and enriched MAIT-cells, 1-2×10^5^ cells were used. For the transwell assay, a 0.3 μm 96 well transwell plate (Corning) was used. 2×10^5^ enriched MAIT-cells were added to the bottom chamber, and 10^6^ PBMCs were added to the top chamber. IFN-α2A (Sigma-Aldrich), IL-12p70 (R&D Systems), IL-18 (R&D Systems), and TNF-α (R&D Systems) were all used at a final concentration of 50 ng/ml. Anti-TNFR2 agonist antibody (clone: MR2-1, Hycult Biotech) was used at a concentration of 2.5 μg/ml. If applicable, inhibitory compounds were added immediately prior to addition of cytokines. Samples were mixed and incubated at 37 °C in 5% CO_2_.

For measurements of MAIT and Vδ2+ T cell activation, Brefeldin A (final concentration of 5 μg/ml; BioLegend) was added after 20 h, and samples were collected after an additional 4 h incubation (24 h total stimulation time). For experiments where cytokine secretion or characteristics of cell transduction were assessed, Brefeldin A was not added and samples were collected after 24 h.

#### Blocking and inhibitory reagents

The following reagents were used in the above-described *in* vitro stimulation assays: mouse IgG1 isotype control antibody (Clone: MOPC-21, BioLegend), mouse IgG2a isotype control antibody (clone: MOPC-173, BioLegend), anti-MR1 antibody (clone: 26.5, BioLegend), anti-IL-12p70 antibody (clone: 24910, R&D Systems), anti-IL-15 antibody (clone: 34559, R&D Systems), anti-IL-18 antibody (clone: 125-2H, R&D Systems), anti-IL-18Ra antibody (clone: 70625, R&D Systems), anti-IFNAR2 (clone: MMHAR-2, Merck Chemicals), B18R (eBioscience), mevastatin (Merck Chemicals), CA-074-Me(*32*) (Merck Chemicals), MCC950(*33*) (Sigma-Aldrich), elevated extracellular K^+^ ion concentration(*34*) (KCl, Sigma-Aldrich), Z-YVAD-FMK(*35*) (R&D Systems), vedolizumab (anti-α4β7 integrin antibody; Takeda Pharmaceuticals), adalimumab (anti-TNF-α antibody; AbbVie Inc), etanercept (TNFR2-Fc fusion protein; Pfizer; A kind gift of Dr. Hussein Al-Mossawi, University of Oxford).

#### Quantification of cytokines and chemokines

For quantification of cell culture supernatant, the following ELISA kits were used: Human TNF-α Quantikine ELISA kit (R&D Systems), IL-18 Human ELISA Kit (MBL International), Human IL-12p70 Quantikine ELISA kit (R&D Systems), and VeriKine Human IFN-α Multi-Subtype ELISA Kit (PBL Assay Science) as per the manufacturer’s instructions. All data were collected on a FLUOstar OPTIMA plate reader (BMG LABTECH). For multiplex analysis of IFN-α-stimulated monocytes the Proteome Profiler Human XL Cytokine Array Kit (R&D Systems) was used as per the manufacturer’s instructions, and the membrane was developed for 10 minutes on an SRX-101A Film Processor (Konica Corporation). For analysis of plasma cytokines in vaccinated human volunteers, the LEGENDplex Human Inflammation Panel (13-plex) (BioLegend) was used per the manufacturer’s instructions. For all assays, samples were diluted as appropriate to fall within the dynamic range of the assay. Samples were freeze-thawed a maximum of one time.

#### MR1 and CD1d tetramer staining in mice and humans

Human and murine MR1/5-OP-RU and MR1/6-FP, and murine CD1d/PBS-57 and CD1d/unloaded tetramers were provided by the NIH Tetramer Facility (Emory University). Tetramers were generated using Phycoerythrin (PE)-Streptavidin and Brilliant Violet 421-Streptavidin (BioLegend) following the NIH Tetramer Facility’s guidelines. All tetramer staining was performed for 40 min at room temperature, and staining was performed in 50 μl of FACS buffer (PBS + 0.05% BSA + 1% EDTA). Following tetramer staining, samples were washed twice in FACS buffer, and used for further staining described below.

#### Surface and intracellular flow cytometry staining of human PBMCs

For activation of MAIT and Vδ2+ T cells, if MR1 tetramer staining was performed, it was done as above. Surface staining was performed using fixable live/dead vital dye (Life Technologies) for 15 min at 4 °C. After surface staining, samples were washed two times in FACS buffer, and cells were fixed and permeabilized for 20 min at 4 °C using Cytofix/Cytoperm (BD Biosciences). Samples were subsequently washed two times in Perm/Wash buffer (BD Biosciences). Intracellular staining was performed for 30 min at 4 °C. The following antibodies were used in Perm/Wash buffer: anti-CD161 (clone: 191B8, Miltenyi Biotec), -CD69 (clone: FN50, BioLegend), -Vα7.2 TCR (clone: 3C10, BioLegend), -Vδ2 TCR (clone: B6, BioLegend), -CD3ε (clone: OKT3 or UCHT1, BioLegend or BD Biosciences), -IFN-γ (clone: B27, BioLegend and BD Biosciences), and - granzyme B (clone: GB11, BD Biosciences). Following intracellular staining, samples were washed two additional times in Perm/Wash buffer, and stored in FACS buffer at 4 °C until analysed on the Flow Cytometer.

For characterization of transduced cell populations, samples were washed two times in FACS buffer, and surface staining was performed for 30 min at 4 °C. The following antibodies were used in FACS buffer: anti-CD11c (clone: B-Ly6, BD Biosciences), -CD19 (clone: HIB19, BioLegend), -CD16 (clone: 3G8, BioLegend and BD Biosciences), -HLA-DR (clone: G46-6, BioLegend), -CD123 (clone: 6H6, BioLegend), -CD14 (clone: M5E2, BioLegend), -CD3ε (clone: UCHT1, BioLegend), and -CD56 (NCAM16.2, BioLegend). After surface staining, samples were washed two times in FACS buffer, and cells were fixed and permeabilized for 20 min at 4 °C using Cytofix/Cytoperm (BD Biosciences). Samples were washed two additional times in FACS buffer and stored at 4 °C until analysed on the flow cytometer. If applicable, intracellular staining for IFN-α2 (clone: 7N4-1, BD Biosciences) was performed following the fixation step. Samples were washed two times in Perm/Wash buffer, and stained in Perm/Wash buffer for 30 min at 4 °C. Following intracellular staining, samples were washed two additional times in Perm/Wash buffer, and stored in FACS buffer at 4 °C until analysed on the flow cytometer.

#### Characterization of murine MAIT and iNKT cells

Following tetramer staining (above), surface staining was performed in FACS buffer for 30 min at 4 °C. The following antibodies were used: anti-CD69 (clone: H1.2F3), -B220 (clone: RA3-6B2), -F4/80 (clone: BM8), -CD44 (clone: IM7), and -TCRβ (clone: H57-597), and fixable live/dead vital dye. All antibodies purchased from BioLegend and vital dye was from Life Technologies. After surface staining, samples were washed two times in FACS buffer, and cells were fixed and permeabilized for 20 min at 4 °C using Cytofix/Cytoperm (BD Biosciences). Cells were subsequently washed two times in Perm/Wash buffer (BD Biosciences). Intracellular staining was performed for 30 min at 4 °C and anti-granzyme B (clone: GB11, BD Biosciences) was added in Perm/Wash buffer. Following intracellular staining, samples were washed two additional times in Perm/Wash buffer, and stored in FACS buffer at 4 °C until analysed on the flow cytometer.

#### Peptide stimulation and intracellular cytokine staining of mouse splenocytes

Peptide restimulation of mouse splenocytes was performed as previously described(*30*). Briefly, 15-mer peptides of the HCV genotype 1b (overlapping by 11 amino acids) were used at a final concentration of 1 μg/ml to stimulate splenocytes for 5 h at 37 °C in 5% CO_2_. Brefeldin A (final concentration of 5 μg/ml; BioLegend) and anti-CD107a antibody (clone: 1D4B, BioLegend) were added at the time of peptide addition. Following stimulation, cells were washed one time in FACS buffer and surface staining was performed for 30 min at 4 °C. The following antibodies were used: anti-CD8a (clone: 53-6.7), -CD4 (clone: RM4-5), -CD44 (clone: IM7), - CD127 (clone: A7R34), and -KLRG1 (clone: 2F1), and Fixable Live/Dead vital dye. All antibodies purchased from BioLegend and vital dye was from Life Technologies. After surface staining, samples were washed two times in FACS buffer, and cells were fixed and permeabilized for 20 min at 4 °C using Cytofix/Cytoperm (BD Biosciences). Cells were subsequently washed two times in Perm/Wash buffer (BD Bioscience). Intracellular staining was performed for 30 min at 4 °C and anti-IFN-γ (clone: XMG1.2, BioLegend) and anti-TNF-α (clone: MP6-XT22, BioLegend) were added in Perm/Wash buffer. Following intracellular staining, samples were washed two additional times in Perm/Wash buffer, and stored in FACS buffer at 4 °C until analysed on the flow cytometer.

#### Data analysis and statistics

All flow cytometry data was acquired on a BD Fortessa Flow Cytometer (BD Biosciences) and processed in FlowJo v. 9.9.6 (FlowJo, LLC). For analysis of the Human XL Cytokine Array, the developed film was scanned (Canon C-EXV), images were converted to greyscale, and pixel density was quantified using ImageJ (v. 1.51). Only membrane spots visible to the naked eye were considered to be positive. All data was analyzed in Prism v. 8.0.1 (GraphPad). For analysis of vaccinated human volunteers, a non-parametric paired Wilcoxon Rank Sum Test was used. For analysis of *in vitro* stimulations, unpaired Student T tests were used for comparison of two groups. A repeated-measures oneway ANOVA with a Dunnett correction for multiple comparisons was used, or a mixed-effects model with Dunnett correction for multiple comparisons was used if there were different numbers of data points in each group. For analysis of dose response curves, a test for linear trend was performed. For analysis of *in vivo* mouse data, a one-way ANOVA was performed with a Dunnett correction for multiple comparisons. For all tests, P<0.05 was considered statistically significant, and P<0.1 was considered a trend with the exact P value reported.

### Supplemental Figures and Legends

**Figure S1.**
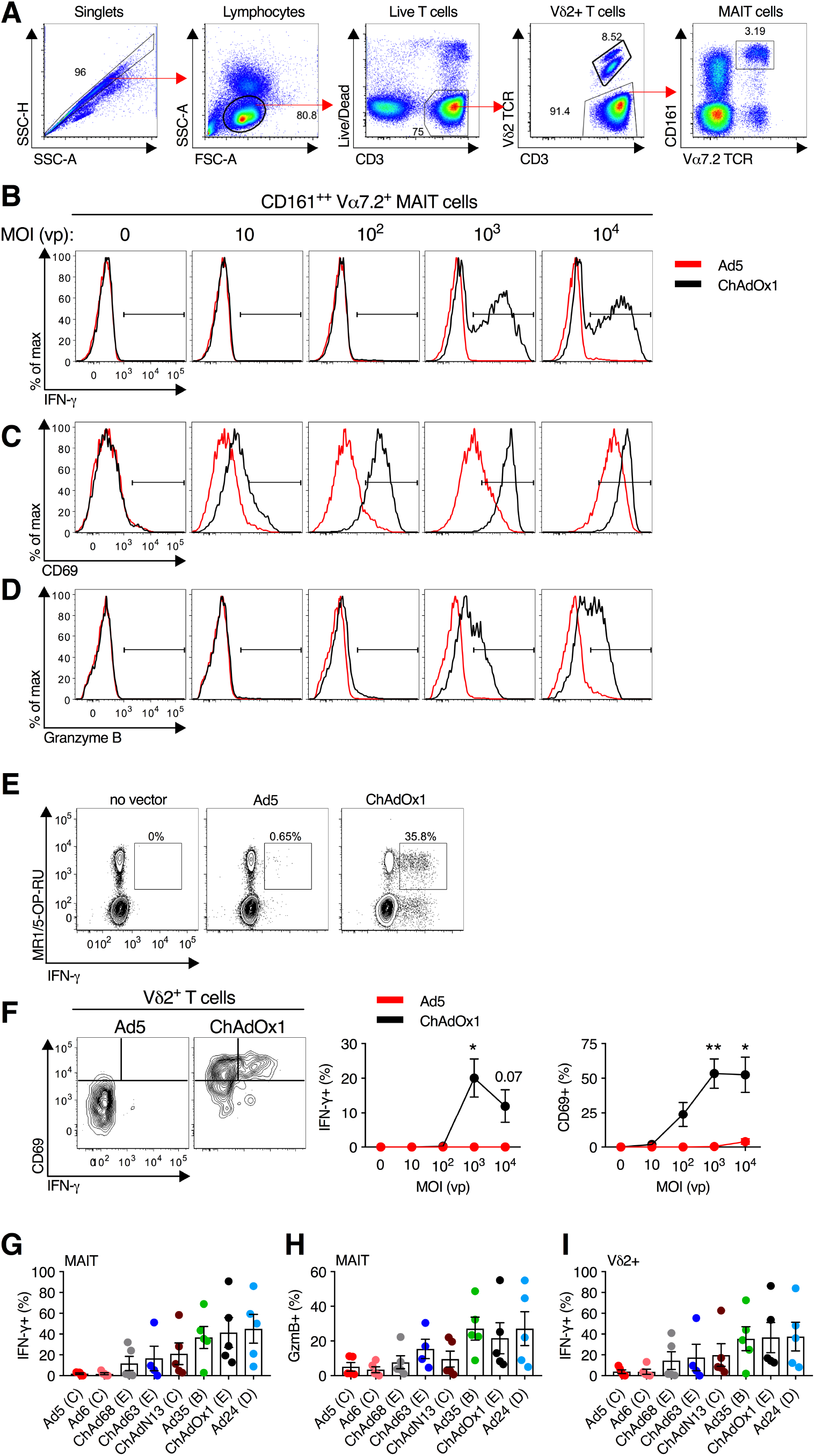
**(A)** Gating scheme for the identification of MAIT cells (CD161++Vα7.2+ T cells) and Vδ2+ T cells in human PBMCs. **(B-D)** PBMCs were stimulated with Ad5 or ChAdOx1 at increasing MOIs (0 to 10^4^ vp), and IFN-γ **(B)**, CD69 **(C)**, and granzyme B (GzmB) **(D)** expression was measured on MAIT cells after 24 h. Representative flow cytometry plots are shown; relates to Fig 1a-c. **(E)** Fresh human PBMCs were stimulated for 24 h with Ad5 or ChAdOx1 (MOI=10^3^ vp), and IFN-γ production by MAIT cells identified using MR1/5-OP-RU tetramers was assessed after 24 h. Data are representative of N=2 donors. **(F)** IFN-γ and CD69 expression on Vδ2+ T cells was measured after 24 h stimulation of PBMCs (N=3) with Ad5 or ChAdOx1 at the indicated MOI. Representative flow cytometry plots (MOI=10^3^ vp) and group averages are shown. **(G-I)** Fresh human PBMCs (N=5) were stimulated with the indicated Ad-vector (species noted in parentheses), and IFN-γ **(G)** or GzmB **(H)** production by MAIT, and IFN-γ production by Vδ2+ T cells **(I)** was measured after 24 h. These data are the individual donor data that relate to Figure 1. *, <0.05; **, P<0.01. Unpaired T test **(F)**. Symbols indicate individual donors, and mean ± SEM are shown.

**Figure S2.**
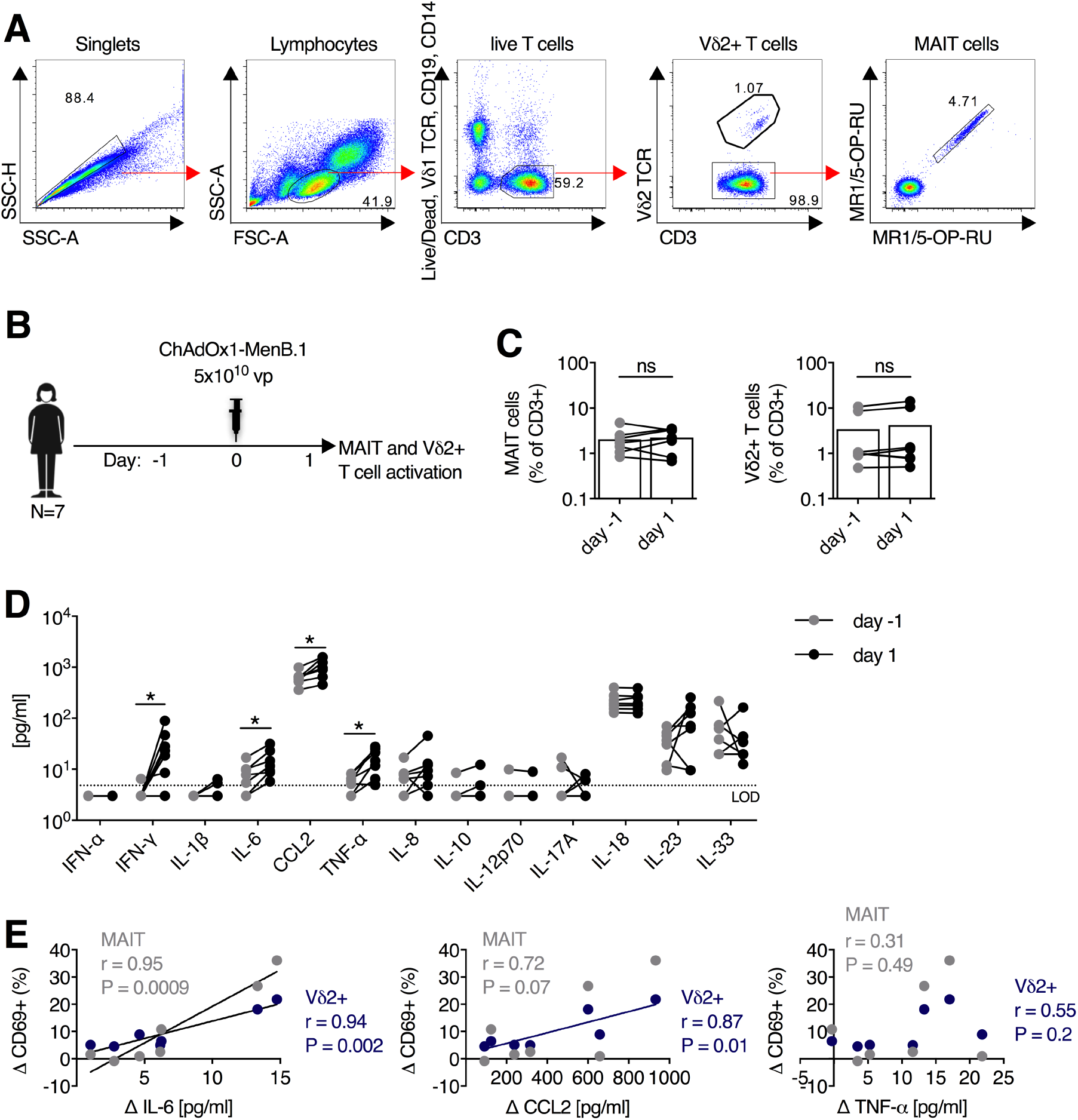
**(A)** Gating scheme for the identification of MAIT cells (MR1/5-OP-RU++ T cells) and Vδ2+ T cells in PBMCs of healthy human volunteers immunized with ChAdOx1. **(B)** Healthy human volunteers (N=7) were immunized with a 5×10^10^ vp dose of ChAdOx1 expressing a *N. meningitidis* group B antigen (MenB.1). **(C)** Frequencies of MAIT cells and Vδ2+ T cells in peripheral blood one day pre- and one day post-immunization. **(D)** Concentration of plasma cytokine levels in healthy human volunteers (N=7) one day pre- and one day post-immunization with 5×10^10^ vp of ChAdOx1-MenB.1. **(E)** Pearson correlation of change in cytokine level following vaccination and the change in expression of CD69 on MAIT cells and Vδ2+ T cells. *, P<0.05; Wilcoxon rank-sum test. Symbols indicate individual donors, and group mean is shown.

**Figure S3.**
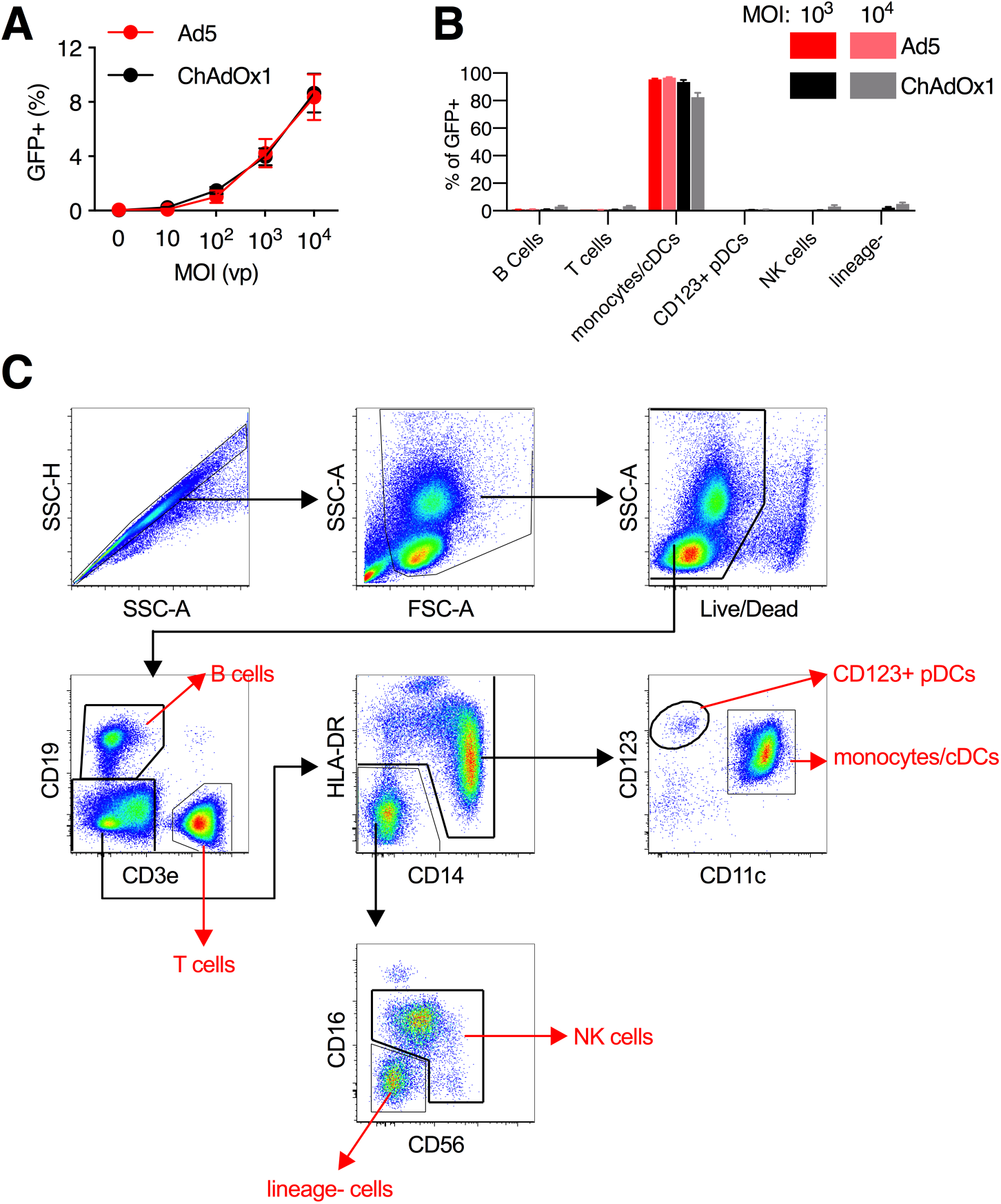
**(A,B)** Fresh human PBMCs were stimulated for 24 h with either Ad5 or ChAdOx1 expressing GFP at the indicated MOI (in vp). **(A)** Fraction of all live PBMCs (N=9) that were GFP+ after 24 h. **(B)** The fraction of GFP+ cells (N=3 donors) that were B cells (CD19+), T cells (CD3+), Monocytes/cDCs (HLA-DR+ CD11c+ CD19- CD3-), CD123+ pDCs (CD123+ CD11c- HLA-DR+ CD3- CD 19-), NK cells (CD56+ CD3-), or lineage- cells (HLA-DR- CD19- CD3- CD56-) was enumerated. **(C)** Gating scheme for the identification of major immune cell subsets within human PBMCs.

**Figure S4.**
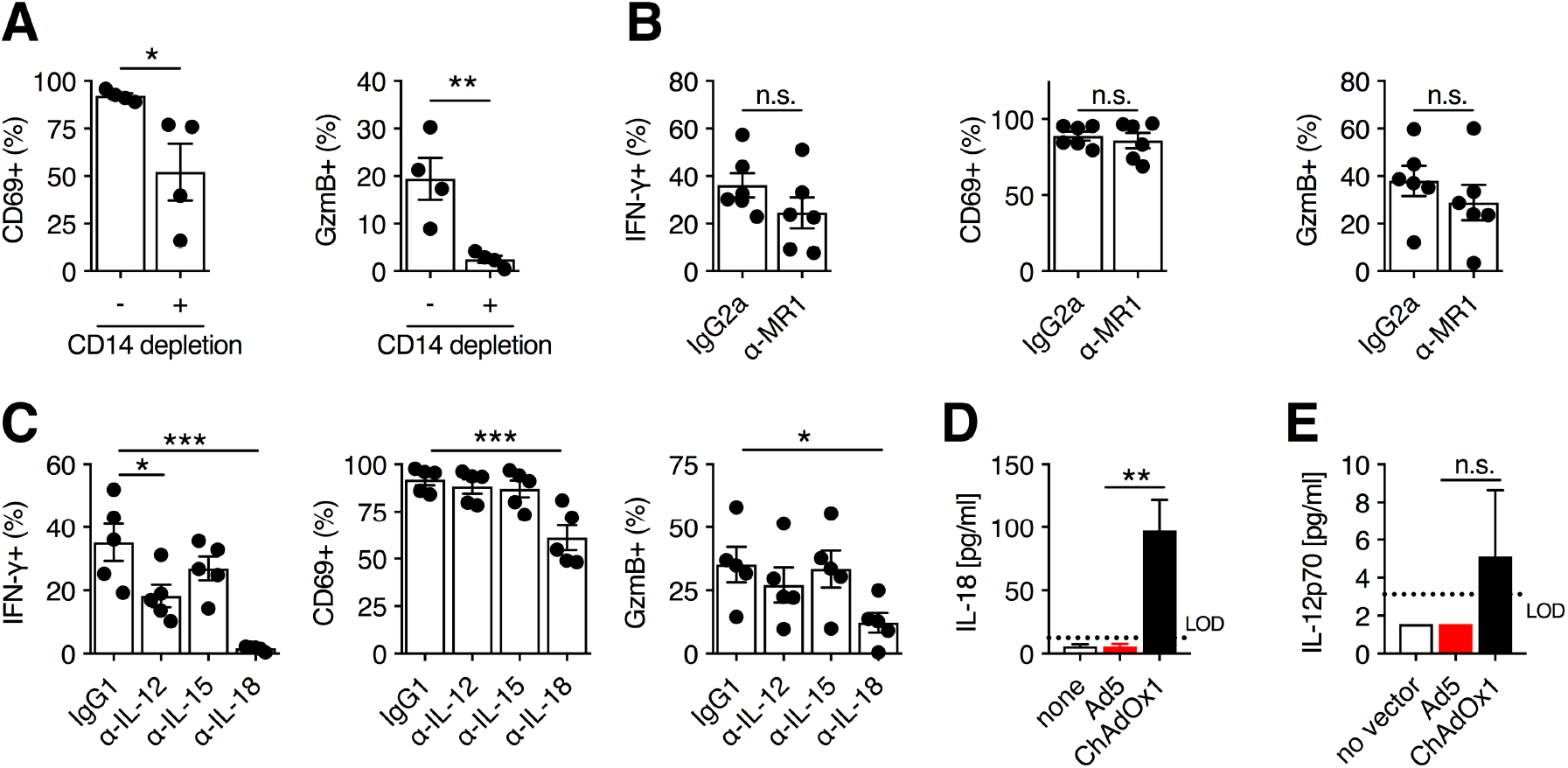
**(A)** PBMCs were depleted of CD14+ monocytes or left untreated as a control (N=4), and were stimulated with ChAdOx1 (MOI= 10^3^ vp). CD69 and granzyme B (GzmB) expression was measured on MAIT cells (CD161++Vα7.2+ T cells) after 24 h. **(B)** Fresh human PBMCs were treated with anti-MR1 antibody (10 μg/ml, N=6) immediately prior to stimulation with ChAdOx1 vectors (MOI=10^3^ vp), and IFN-γ, CD69, and GzmB expression on MAIT cells was measured after 24 h. **(C)** PBMCs (N=5) were stimulated with ChAdOx1 (MOI=10^3^ vp), and anti-IL-12, anti-IL-15, or anti-IL-18 antibodies (10 μg/ml) were added immediately prior to vector addition. IFN-γ, CD69, and GzmB expression was measured on MAIT cells after 24 h. Concentration of IL-18 **(D)** and IL-12p70 **(E)** in cell culture supernatants following 24 h stimulation of fresh PBMCs with Ad5 or ChAdOx1 (MOI=10^3^ vp). *, P<0.05; **, P<0.01; ***, P<0.001. Unpaired T test **(A,B,D,E)** and repeated-measures one-way ANOVA with Dunnett correction **(C)**. Symbols indicate individual donors, and mean ± SEM are shown.

**Figure S5.**
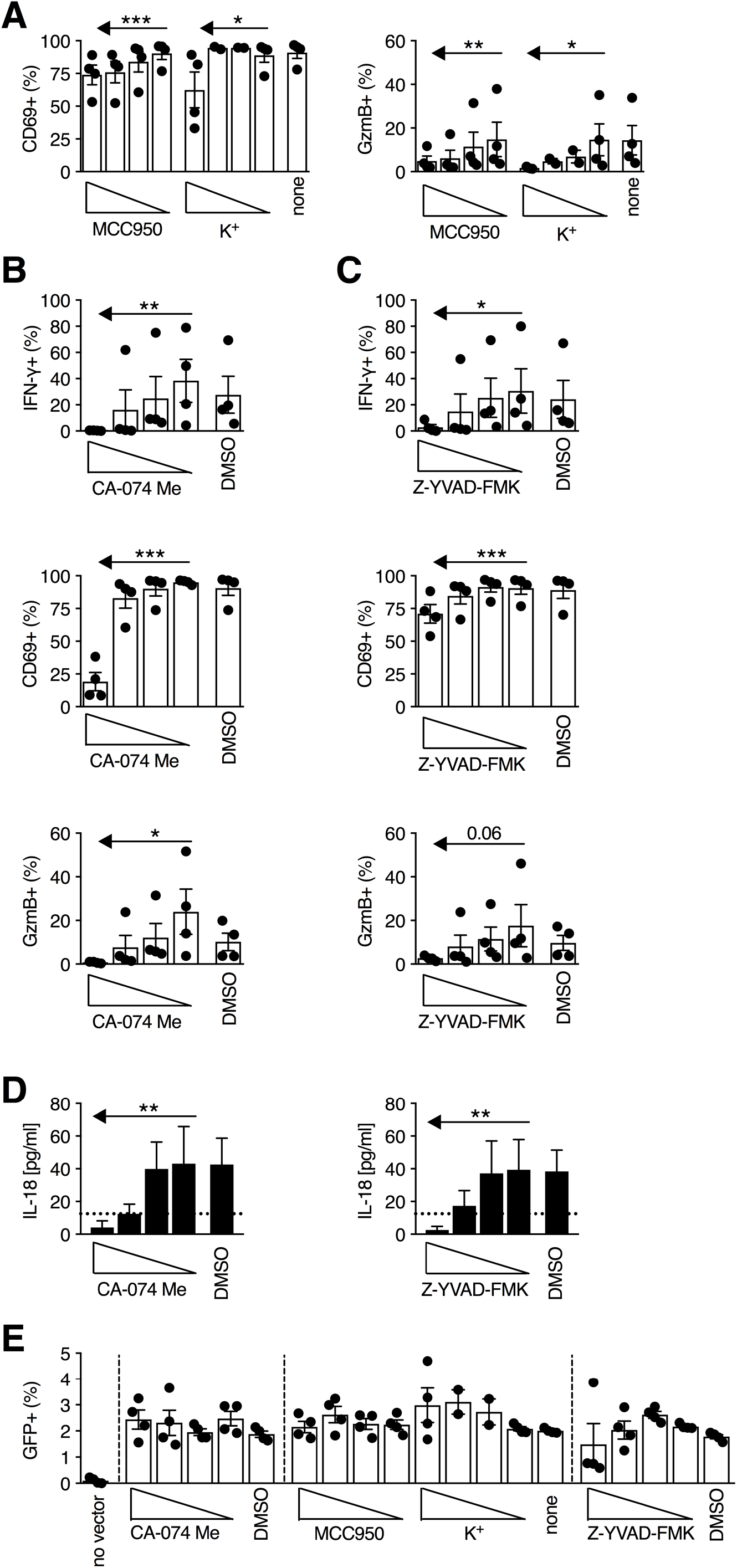
**(A)** The NLPR3 inhibitors (MCC950 [0.01-10 μM]) and extracellular K+ [5-30 mM]) were added immediately prior to stimulation of PBMCs with ChAdOx1 (MOI=10^3^ vp), and CD69 and granzyme B (GzmB) production by MAIT cells (CD161++Vα7.2+ T cells) was assessed after 24 h (N=4). Cathepsin B inhibitor (CA-074-Me [0.01-10 μM]) **(B)** and Caspase 1 inhibitor (Z-YVAD-FMK [0.1-100 μM]) **(C)** were added immediately prior to stimulation of PBMCs with ChAdOx1 (MOI=10^3^ vp). After 24 h, IFN-γ, CD69, and GzmB production by MAIT cells was assessed (N=4). **(D)** Concentration of IL-18 in cell culture supernatants of PBMCs treated with the indicated inhibitors 24 h after stimulation with ChAdOx1 MOI=10^3^ vp; N=4). **(E)** Frequency of GFP+ PBMCs (N=4) at 24 h following transduction with ChAdOx1 expressing GFP (MOI=10^3^ vp) in the presence of the indicated dose of CA-074 Me, MCC950, extracellular K+ ions, or Z-YVAD-FMK. *, P<0.05; **, P<0.01; ***, P<0.001. Repeated-measures one-way ANOVA with test for linear trend. Symbols indicate individual donors, and mean ± SEM are shown.

**Figure S6.**
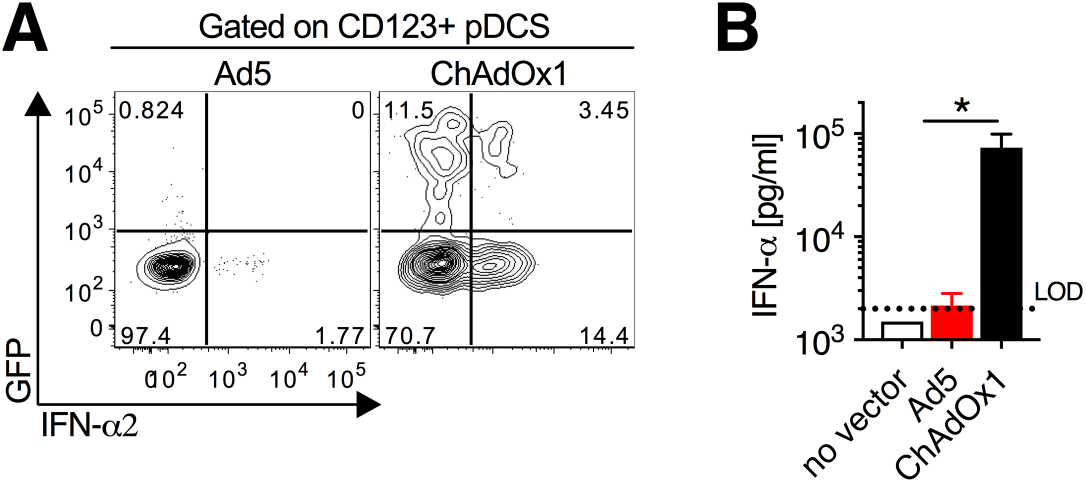
**(A)** Expression of IFN-α2 on CD123+ pDCs (HLA-DR+ CD11c- CD14- CD16- CD19- CD3-) 24 h after stimulation with Ad5 or ChAdOx1 expressing GFP (MOI=10^3^ vp). Data are representative of N=4 donors. **(B)** Concentration of IFN-α in cell culture supernatants following 24 h stimulation of fresh PBMCs with Ad5 or ChAdOx1 (MOI=10^3^ vp). *, P<0.05. Unpaired T test **(B)**. Mean ± SEM are shown.

**Figure S7.**
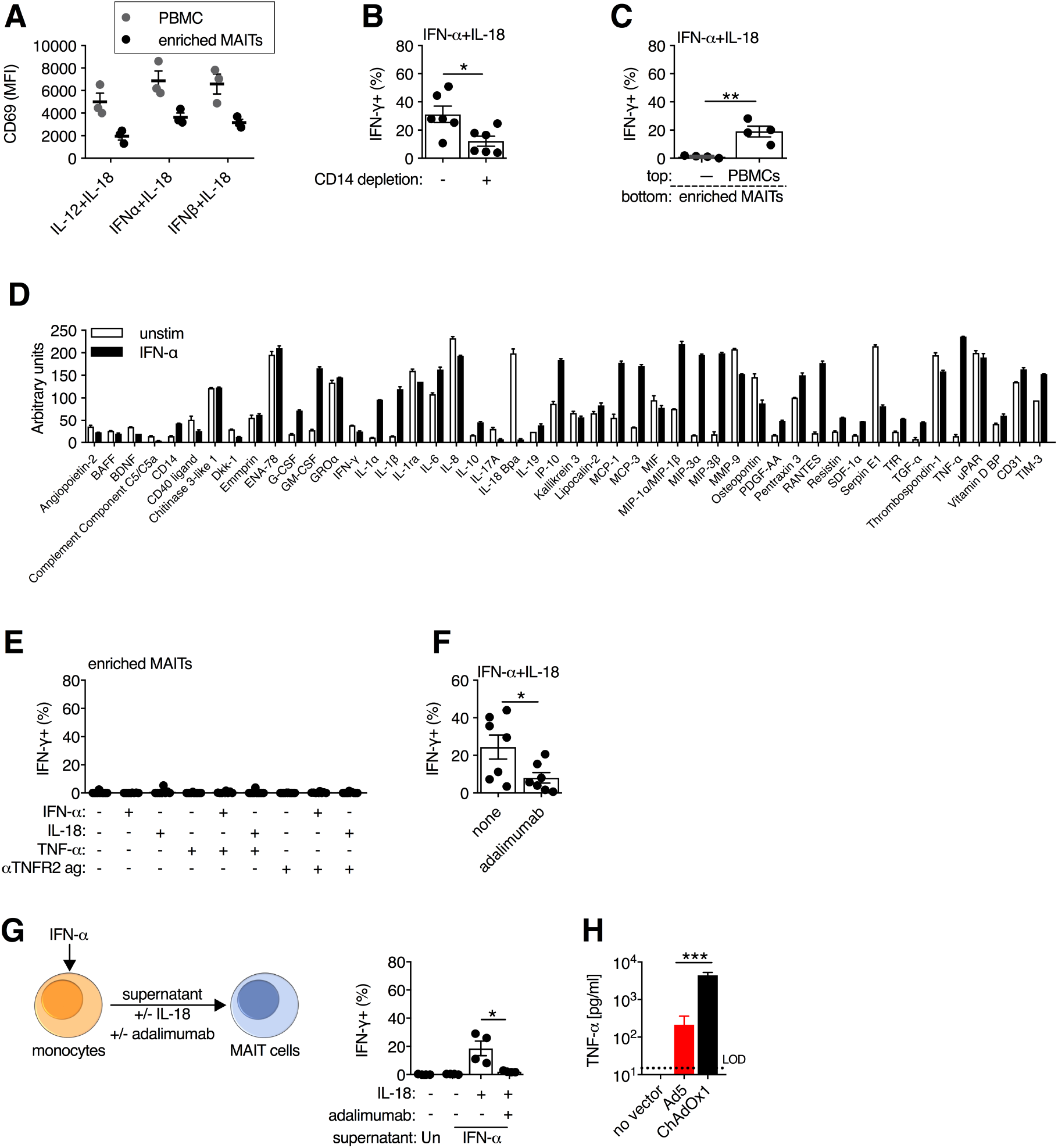
**(A)** Unfractionated PBMCs or enriched MAIT cells (positive selection by CD8 MicroBeads) were stimulated for 24 h with the indicated cytokines (50 ng/ml), and CD69 expression was measured on MAIT cells after 24 h. **(B)** Unfractionated PBMCs or CD14+ monocyte-depleted PBMCs were stimulated with IFN-α + IL-18 (50 ng/ml), and IFN-γ production by enriched MAIT cells was measured after 24 h (N=6). **(C)** Using a 0.3 μm transwell system, enriched MAIT cells were placed in the bottom chamber and the top chamber was loaded with either autologous PBMCs or left cell-free (N=4). IFN-γ production by MAIT cells was assessed at 24 h following stimulation with IFN-α + IL-18 (50 ng/ml). **(D)** CD14-purified monocytes (N=1, in duplicate) were stimulated with IFN-α (50 ng/ml) or left untreated. After 24 h, supernatants were collected and used fresh for immunoblotting. Relative protein concentration was calculated by quantifying pixel density. **(E)** CD8- MicroBead enriched MAIT cells (N=10) were stimulated with single or double combinations of IFN-α, IL-18, TNF-α, or anti-TNFR2 agonist antibody (cytokine at 50 ng/ml and agonist antibody at 2.5 μg/ml). IFN-γ production by MAIT cells was measured after 24 h. **(F)** PBMCs (N=7) were stimulated with IFN-α + IL-18 (50 ng/ml), and adalimumab (anti-TNF-α antibody; 10 μg/ml) was added immediately prior cytokine addition. IFN-γ production by MAIT cells was measured after 24 h. **(G)** Purified monocytes (N=4) were stimulated with IFN-α (50 ng/ml), or left untreated, and after 24 h supernatants were transferred to autologous enriched MAIT cells ± IL-18 (50 ng/ml) and ± adalimumab (anti-TNF-α antibody; 10 μg/ml). IFN-γ production by MAIT cells was measured at 24 h. **(H)** Concentration of TNF-α in cell culture supernatants of PBMCs 24 h after stimulation with Ad5 or ChAdOx1 (MOI=10^3^ vp; N=7). *, P<0.05; **, P<0.01; ***, P<0.001. Unpaired T test. Symbols indicate individual donors, and mean ± SEM are shown.

**Figure S8.**
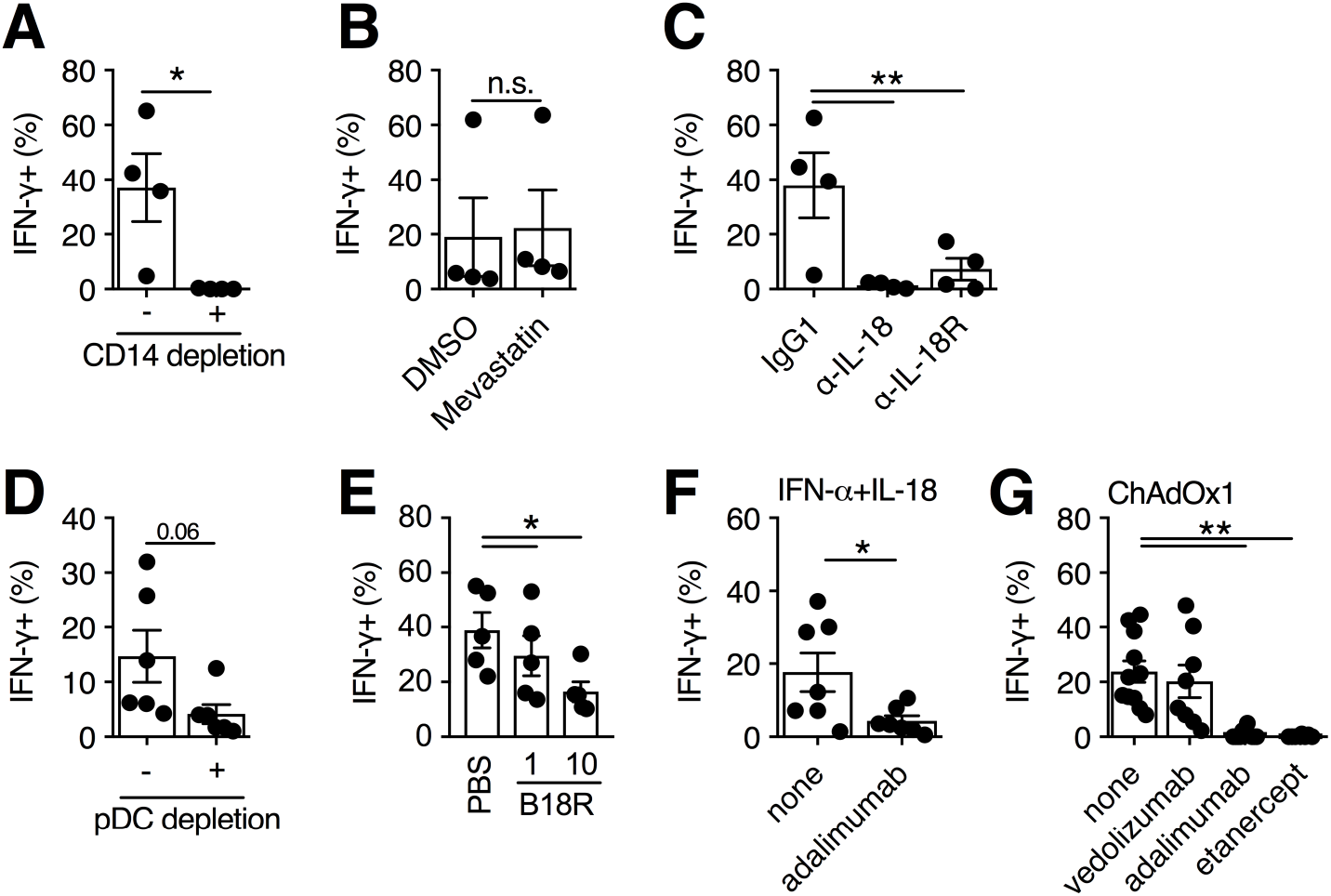
**(A)** Fresh human PBMCs were depleted of CD14+ monocytes or left untreated as a control (N=4), and were stimulated with ChAdOx1 (MOI=10^3^ vp). IFN-γ production by Vδ2+ T cells was measured after 24 h. **(B,C)** Fresh human PBMCs were treated with mevastatin (5 μM; N=4) (B), or either anti-IL-18 or anti-IL-18R antibodies (10 μg/ml; N=4) **(C)** immediately prior to addition of ChAdOx1 (MOI=10^3^ vp). IFN-γ production by Vδ2+ T cells was measured after 24 h. **(D)** PBMCs were depleted of CD123+ pDCs or left untreated as a control (N=4), and IFN-γ expression was measured on Vδ2+ T cells after 24 h stimulation with ChAdOx1 (MOI=10^3^ vp). **(E)** PBMCs (N=5) were stimulated with ChAdOx1 (MOI=10^3^ vp), and B18R (1 or 10 μg/ml) was added immediately prior to vector addition. IFN-γ production by Vδ2+ T cells was measured after 24 h. **(F)** PBMCs (N=7) were stimulated with IFN-α + IL-18 (50 ng/ml), and adalimumab (anti-TNF-α antibody; 10 μg/ml) was added immediately prior to cytokine addition. IFN-γ production by Vδ2+ T cells was measured after 24 h. **(G)** PBMCs were stimulated with ChAdOx1 (MOI=10^3^ vp), and vedolizumab (anti-α4β7 integrin antibody, N=8), adalimumab (anti-TNF-α antibody, N=11), or etanercept (TNFR2-Fc fusion protein, N=8) (10 μg/ml) was added immediately prior to vector addition. IFN-γ production by Vδ2+ T cells was measured after 24 h. *, P<0.05; **, P<0.01. Unpaired T test. **(A,B,D,F)** and repeated-measures one-way ANOVA with Dunnett Correction **(C,E,G)**. Symbols indicate individual donors, and mean ± SEM are shown.

**Figure S9.**
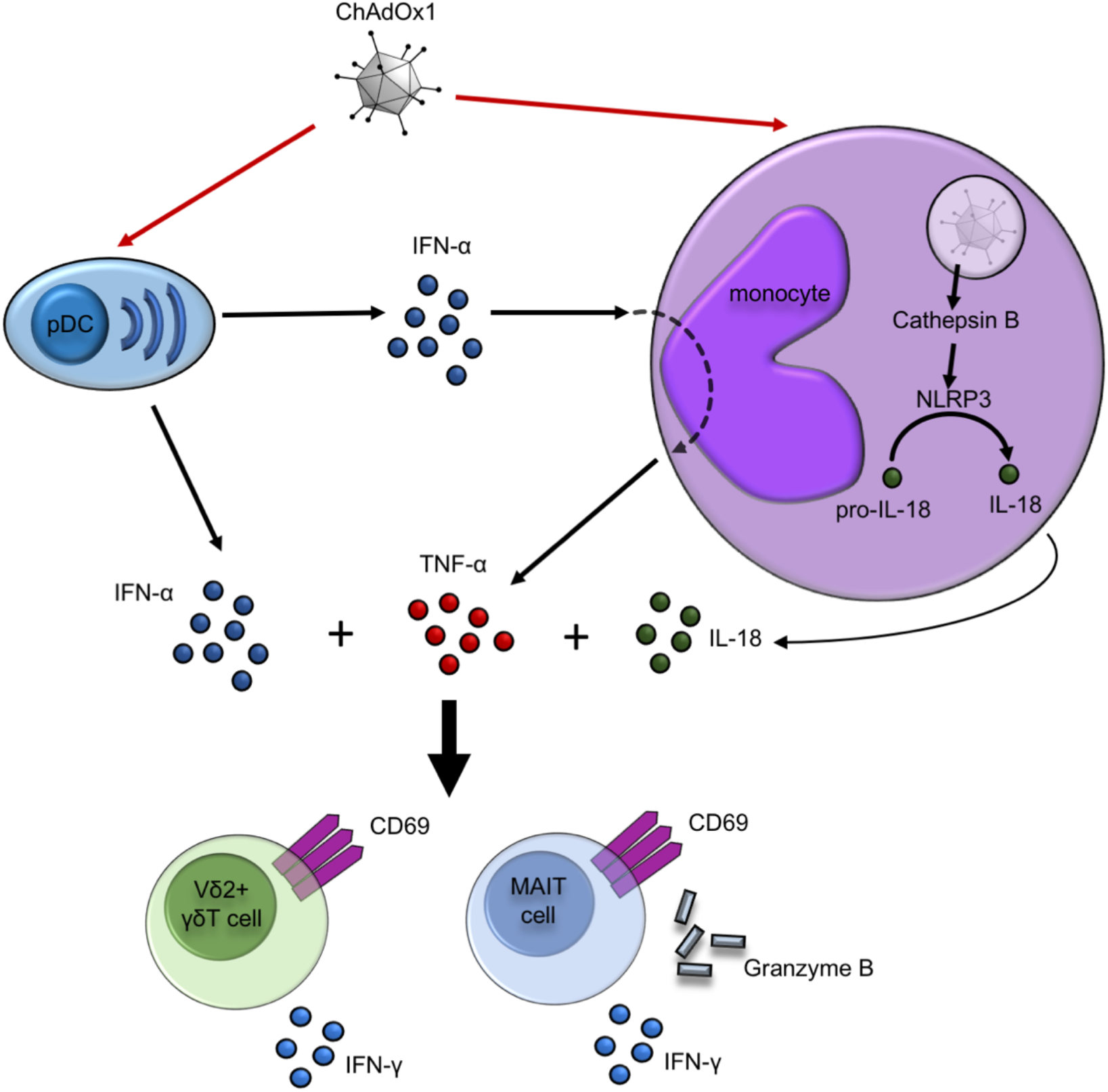
Model for how ChAdOx1 (and other stimulatory Ad vectors) activates MAIT cells and Vδ2+ T cells. Activation involves two pathways: (1) ChAdOx1 transduces monocytes and induces IL-18 production by activating the NLRP3 inflammasome, and (2) ChAdOx1 transduces pDCs and triggers the production of IFN-α, which in turn drives TNF-α production by monocytes. The combination of IL-18, IFN-α, and TNF-α act on MAIT and Vδ2+ T cells to induce expression of IFN-γ, Granzyme B, and CD69.

**Figure S10.**
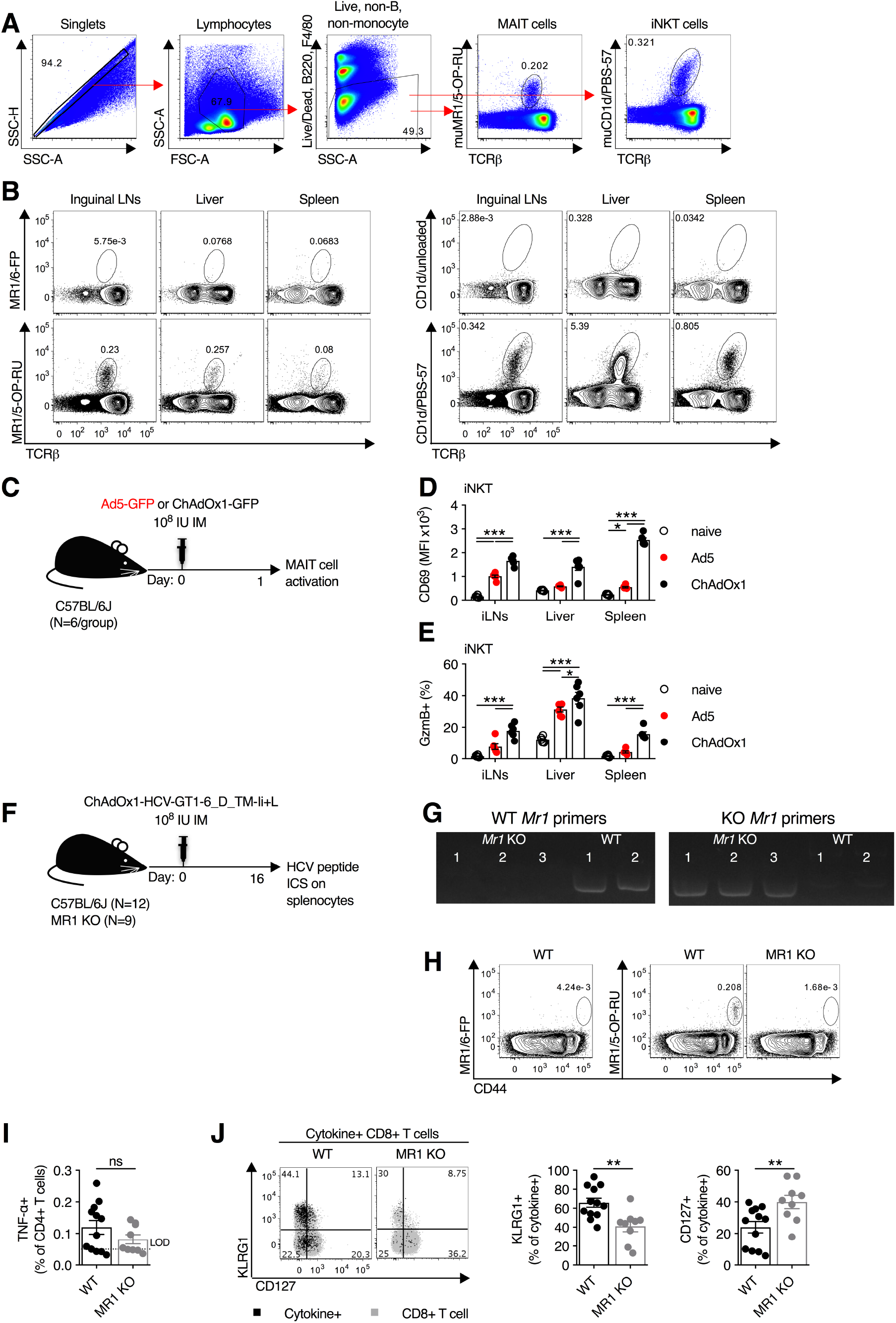
**(A)** Gating scheme for the identification of MAIT cells and iNKT cells in mice using MR1 and CD1d tetramers. **(B)** Representative flow cytometry plots of the frequency of MAIT cells and iNKT cells in the inguinal LNs, liver, and spleen of C57BL/6 mice. **(C-E)** C57BL/6J mice (N=6 per group) were immunized intramuscularly with 10^8^ IU of Ad5 or ChAdOx1 expressing GFP (C), and one day post-immunization expression of CD69 **(D)** and granzyme B (GzmB) **(E)** was measured on iNKT cells (CD1d/PBS-57+ T cells) isolated from inguinal LNs, liver, and spleen. Data are representative of two independent experiments. **(F-J)** C57BL/6J (N=12) or MR1 KO (N=9) mice were immunized intramuscularly with 10^8^ IU of ChAdOx1 expressing HCV-GT1-6_D_TM-Ii+L transgene, and on day 16 post-immunization splenocytes were collected **(F)**. **(G,H)** Genotyping of wild-type C57BL/6 and MR1 KO mice by PCR for the *Mr1* gene **(G)** and confirmation of the absence of MAIT cells in the inguinal LNs by flow cytometry **(H)**. **(I)** Group averages for TNF-α production by CD4 T cells following 5 h restimulation with an overlapping peptide pool of the HCV genotype 1b proteome. **(J)** Representative flow cytometry plots and group averages of KLRG1 and CD127 expression on cytokine-producing CD8 T cells following peptide restimulation. *, P<0.05; **, P<0.01; ***, P<0.001. One-way ANOVA with Sidak correction for multiple comparisons **(D,E)** and unpaired T test **(I,J)**. Symbols indicate individual animals, and mean ± SEM are shown.

